# Dopamine Agonists in Parkinson’s Disease Decouple Risk-Taking from Reward-Paired Cues by Enhancing Risky Choice in Their Absence

**DOI:** 10.64898/2026.09.05.748022

**Authors:** Katherine Muksuris, Ellen Flynn, Tanya L. Feng, Pratibha Surathi, Shanna C. Yeung, Leili Mortazavi, Anna Isberg, Martin J. McKeown, Silke Appel-Cresswell, Catharine A. Winstanley, A. Jon Stoessl, Jason J.S. Barton, Mariya V. Cherkasova

## Abstract

**Background:** Common side effects of dopamine replacement therapy (DRT) for Parkinson’s Disease (PD) are addiction-like impulse control disorders, including compulsive gambling. Gambling products such as slot machines prominently feature reward-paired audiovisual stimuli, which can promote riskier choices on laboratory gambling tasks in both humans and rodents. In rats, risky decision making in the presence of reward-paired cues is dependent on dopamine D3 receptor activity and is increased by ropinirole, a D2/3 receptor agonist with a higher affinity for D3 receptors. In humans, it remains uncertain whether cue-induced risky choice is amplified by dopamine agonism.

**Methods:** We tested effects of DRT on cue-induced risky choice in 62 patients with PD (34 on levodopa monotherapy, 27 on levodopa and dopamine agonists). Patients performed two versions of a risky decision-making task: one with and one without reward-paired sensory cues, both ON and OFF DRT. A group of 36 age-matched controls completed the same task versions without DRT.

**Results:** Across all groups, participants made riskier choices when the task included reward-paired sensory cues. Levodopa monotherapy did not affect risky decision-making. However, a combination of levodopa and dopamine agonists increased risky choices specifically in the uncued version of the task, making performance indistinguishable from that on the cued task.

**Conclusions:** Dopamine agonists combined with levodopa promote risk-taking even in the absence of reward-paired cues. This could translate into risky reward-seeking behaviors that are decoupled from contextual factors.

## INTRODUCTION

Dopamine replacement therapy (DRT) is the mainstay treatment for Parkinson’s disease (PD). Its two main forms are levodopa (co-administered with carbidopa to prevent peripheral decarboxylation to dopamine) and dopamine receptor agonists (DAs). Treatment with DRT, particularly DAs, significantly increases the risk of developing impulse control disorders (ICDs), one of which is gambling disorder (1–6). ICDs usually resolve when the DRT is reduced in dose or discontinued (4,5,7) but may persist (8). Neurobehavioral mechanisms by which DRT produces iatrogenic gambling disorder and other ICDs are not fully understood. One hypothesized mechanism is the effect of DRT on risky decision making, with increased dopamine signaling proposed to encourage risky behaviors. Whereas unmedicated patients with PD are generally risk-averse (9–11), patients on DRT are more likely to make risky decisions on laboratory tasks compared to unmedicated patients and controls (4,12–15), although results have been mixed (16–19).

In people without PD, problematic gambling is thought to result from person-level vulnerabilities (e.g. genetic and neurobiological predisposition) interacting with characteristics of gambling products and environments (20,21). A prominent feature of gambling products are salient audiovisual stimuli that often accompany, signal or foreshadow winning outcomes. Although the role of these audiovisual reward cues in promoting problematic gambling is incompletely understood (22–25), evidence in rats and humans suggests that they promote riskier decision making on laboratory gambling tasks (26–33). In rats, risk-taking propensity is associated with increasing preference for cued versus ‘uncued’ gambling tasks over time (34). Furthermore, acute dopamine D3 receptor agonism enhanced, while antagonism neutralized, the risk-promoting effect of reward-paired cues on the rodent gambling task (35). Analogously, chronic administration of ropinirole, a D2/3 agonist commonly used in the treatment of PD, during rodent gambling task acquisition resulted in risk-preferring performance if the task included reward-paired cues, but not on the ‘uncued’ task (32).

While this evidence implicates D3 receptor signaling in the risk-promoting effects of reward-paired cues, this is currently limited to animal literature. The possible role of DAs in the risk-promoting effects of reward-paired cues in humans remains unknown, as are the possible contributions of these effects to the iatrogenic gambling disorder in PD. The current study aimed to investigate those effects. We tested effects of levodopa monotherapy versus dual therapy with levodopa and DAs, hypothesizing that both types of DRT, and especially dual therapy, would enhance risk-promoting effects of reward-paired cues in a laboratory gambling task.

## METHODS AND MATERIALS

### Participants

We tested 61 patients with idiopathic PD. Thirty-four were receiving levodopa monotherapy (LD) and 27 a combination of levodopa and DAs (LD+DA). Thirty-six non-PD Controls were also tested. Controls were age-matched to patients within 5 years; gender was matched except in two cases where the opposite sex spouse participated. All patients were recruited from the Movement Disorders Clinic at the University of British Columbia and provided written informed consent prior to participation. The study was conducted in accordance with institutional guidelines and the Declaration of Helsinki and was approved by the Research Ethics Board of the University of British Columbia.

Inclusion criteria were age ≥ 18 years, normal or corrected-to-normal vision and hearing, and, for the patients, treatment with a stable dose of oral DRT (LD or LD+DA) and ability to withhold DRT for ≥ 12 hours without complications. Treatment with deep brain stimulation or infusion device was exclusionary. Other exclusion criteria were a Montreal Cognitive Assessment (MoCA) (36) score < 20, other neurological diagnoses, or evidence of past or present ICDs based on the patient’s chart or in-session questionnaires (see below). One patient in the LD group was excluded from analyses due to a MoCA score of 17.

### Procedure

Patients were tested both ON and OFF their DRT in two separate sessions, with session order (ON or OFF) randomized across participants. Controls were tested twice without DRT, with the task versions (see below) in both sessions corresponding to that for the matched patient. The two sessions were held 6 days apart on average (M= 6.17, SD = 6.11, range: 1-30 days). For the analyses, the ON/OFF order variable was coded for each control participant based on the order of their matched patient, as Controls did not receive DRT. In the OFF session, PD patients were instructed to withhold immediate-release levodopa for 12 hours and controlled release levodopa or DAs for 18 hours prior to testing. In the ON session, patients were tested on their usual DRT.

In each session, participants completed two versions of a risky decision-making Vancouver Gambling Task (VGT, described below): one included audiovisual reward cues (“cued”) while the other did not (“uncued”), with version order randomized across participants. In between the two versions, the Movement Disorders Society Unified Parkinson’s Disease Rating Scale (MDS-UPDRS) Part III (37) was administered (in both sessions, ON and OFF) by a trained research assistant, and video-recorded for blinded scoring. Due to insufficient recording quality, MDS-UPDRS-III could not be rated for two LD and 12 LD+DA participants in the OFF state and for 3 LD and 11 LD+DA participants in the ON state. Following the cued and uncued VGT, two other decision-making tasks (not reported here) and questionnaires (described below) were completed.

### Vancouver Gambling Task

The Vancouver Gambling Task (VGT) (15,26,27,38,39) is a two-alternative behavioral economic task assessing willingness to take risks depending on the probability and magnitude of the expected reward. Our group has previously used this task to assess risky decision making in healthy individuals (26,27,38) and PD (15,39). Participants were presented with two probabilistic monetary prospects: a smaller but more probable gain versus a larger but less probable gain (i.e. the “riskier” alternative). “Risk” is used here to indicate opting for a lower-probability outcome rather than to denote variance in prospective outcome in a classic behavioral-economic sense. Losses were not possible. The probabilities associated with each of the alternatives (Figure 1) always added up to 100% (60/40, 70/30, 80/20). A higher reward probability was always paired with a smaller magnitude and vice versa. Magnitudes ranged from 1 to 5 units, each equating to 10¢ Canadian. Each task version comprised 10 unique pairs of alternatives, each presented 10 times in pseudo-random order for a total of 100 trials. The pairs were defined by their relative expected values (EV), where EV equals the probability multiplied by the magnitude of reward. Hence, each pair can be described using its Expected Value Ratio (EVR): EVR = EV_(safe)_- EV_(risky)_/mean(EV_(safe)_, EV_(risky)_). Following each choice, participants received outcome feedback (a win or a non-win). The total sum won on the tasks was added to participants’ study compensation.

**Figure 1:**
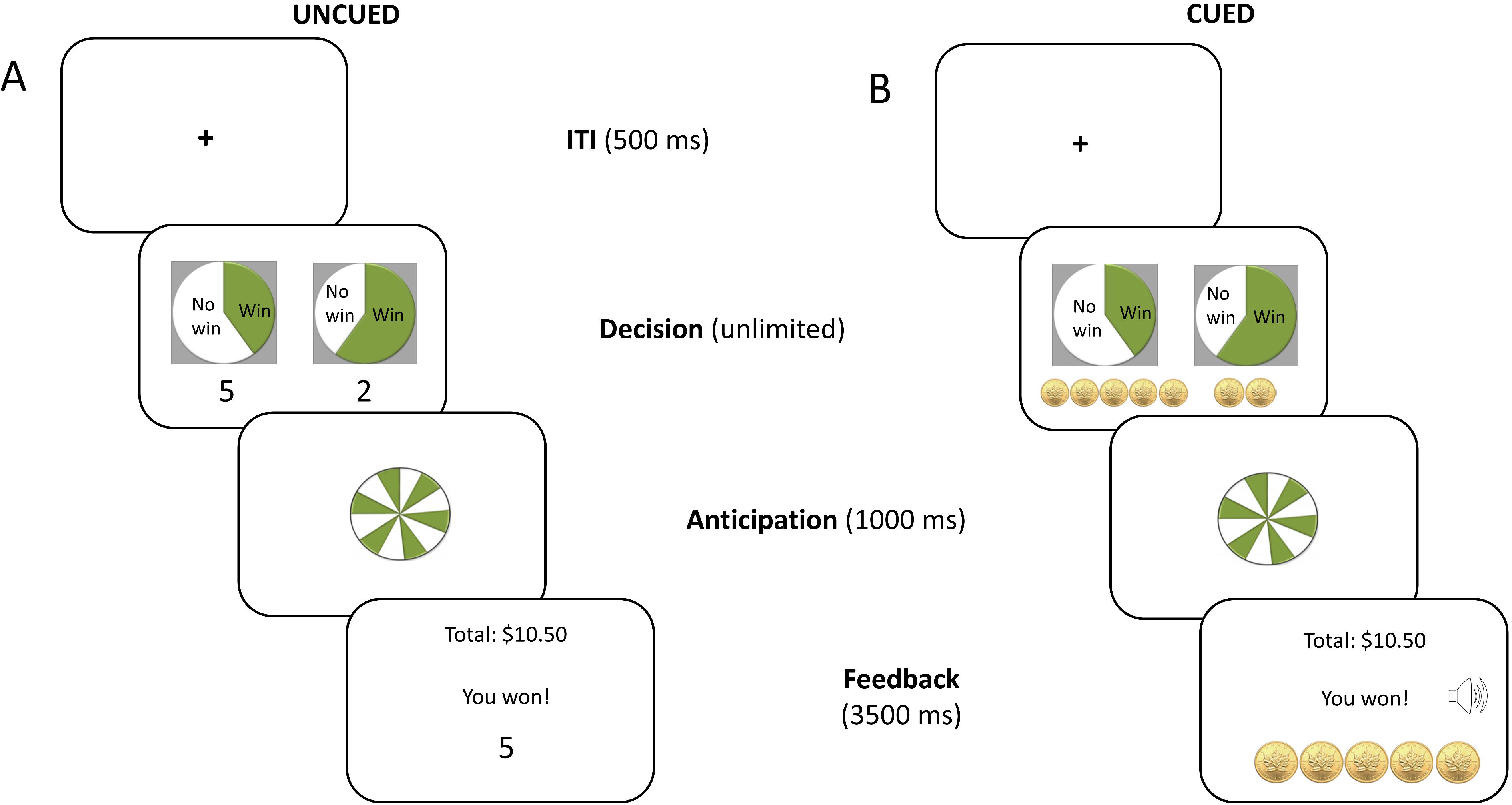
Schematic of the Vancouver Gambling Task in both the A) uncued and B) cued conditions. Pie charts represented probabilities for each option: 20% vs. 80%, 30% vs. 70%, or 40% vs. 60%. The gain magnitudes represented using either numerals (uncued) or gold coins (cued) beneath the pie charts, indicated the number of tokens that could be won (1, 2, 3, 4 or 5), with each token worth ¢10 Canadian in actual money. The position of the riskier (i.e. larger-magnitude) option on the right or the left of the screen was randomized across trials. The decision phase of the task was followed by an anticipation phase with a spinning roulette display. A feedback phase then followed with (cued) or without (uncued) audiovisual cues.

Each session had an uncued VGT version and a cued one. In the uncued version, the magnitudes of potential gains in the choice phase of the task and the winning outcomes were displayed by numbers (Figure 1A). In the cued version, potential reward magnitudes were represented by gold coins in the choice phase. In the feedback phase, outcomes were represented using static or dynamic images of the coins won, accompanied by casino-inspired jingles (Figure 1B). The sensory intensity and complexity (e.g. sparkles, motion, sound length and complexity) scaled with win size (26).

Participants were given five practice trials at the beginning of each version and an optional break after every 20 trials. There were two versions each of the cued and uncued task with slightly different EVs in each pair to mitigate practice effects between the two testing sessions.

VGT was programmed in Experiment Builder and accompanied by eye tracking using EyeLink 1000 (SR Research Ltd., Mississauga, Ontario), not reported here.

### Questionnaires

The MoCA (36) was conducted to ensure capacity to complete the tasks. Mood symptoms were assessed using the Beck Depression Inventory (BDI) (40) and Lille Apathy Rating Scale (LARS) (41). ICD symptoms were assessed using the Questionnaire for Impulsive-Compulsive Disorders in Parkinson’s Disease – Rating Scale (QUIP-RS) (42); the Novelty Seeking Scale of the Temperament and Character Inventory (TCI-NS) (43) was used as a dispositional measure of impulsiveness. Additionally, gambling problems were assessed using the Problem Gambling Severity Index (PGSI) (44).

### Statistical Analyses

Analyses were conducted using R (Version 4.4.3) (45) and GraphPad Prism 10.2 (46). Group demographics, clinical characteristics and questionnaire scores were analyzed using one-way ANOVAs (or the non-parametric equivalent) or 2-sample t-tests for continuous variables and chi-square tests for categorical variables.

Effects of cues and medication on VGT choices were analyzed separately in each group (LD, LD+DA, Controls) with Bayesian mixed effects logistic regression models using the *brms* R package (47). For the Controls, the Medication variable was substituted with a Session variable that took on the matched patients’ Medication variable values to account for effects of repeated testing. Linear mixed effects models with a logistic link using the lme4 package (48) were first attempted, but diagnostics revealed influential observations, so Bayesian models were fit instead as a robust alternative (this did not change the findings).

The choice of a risky (versus safe) option was predicted on a trial-by-trial-basis as a function of EVR in interaction with Cues (cued vs uncued) and Medication (ON vs OFF, or Session for Controls). Random slopes were modeled for EVR and for the interaction of Cues with Medication state, and random intercepts were included for participants. All categorical variables were sum-coded except for the Group variable in the comparison of uncued VGT performance in the OFF state across groups (see below), in which Controls were the reference group. Covariates were selected using a model comparison procedure, in which potential covariates were added one-by-one and retained if they improved model fit based on AIC and BIC criteria. Task Version (gamble EVs and EVRs), testing session Order (ON or OFF first), MDS-UPDRS Part III score in the OFF state (scaled variable), and Trial number (1-100, scaled variable) were included as covariates in the final model in each group (except for MDS-UPDRS-III in the model for Controls). A random slope was also modeled for the Trial number covariate. See Supplement for best-fitting model formulas.

A similar Bayesian model was fit to compare the three groups on risky choice in the OFF state (or corresponding Session) on the uncued version of the VGT to establish whether they differed in risky choice propensity independent of cue and medication effects. Risky choice was modeled as a function of EVR in interaction with Group. Task Version, testing session Order, MDS-UPDRS Part III score in the OFF state, and Trial were included as covariates. Random slopes were modeled for the effects of EVR and Trial, and random intercepts were included for participants. The model formula is below:

Priors for the intercept and the fixed effect of EVR were informed by a previous study of an independent sample of young healthy controls (27). For the intercept, a normal prior with the mean of -0.66 and a standard deviation (SD) of 2.5 was used; for EVR, a normal prior with the mean of 3.0 and an SD of 2.5 was used. For all other fixed effects, weakly informative default student-t priors with the mean of 0 and SD of 2.5 were used. For random effects SD, a weakly informative exponential prior distribution with a mean and SD of 1 was used. For the random effects correlation matrices, the default Lewandowski-Kurowicka-Joe (LKJ) prior with the shape parameter of 1 was used, corresponding to a uniform distribution of the shape of the correlation matrices (49). The models were estimated using MCMC sampling with 4 chains of 4000 iterations and a warmup of 2000. Diagnostics were performed by visual inspection of the caterpillar plot. The convergence of chains was determined by the R-hat statistic, with values between 1 and 1.01 considered acceptable. Statistical significance of coefficients was determined by the 95% Bayesian credible interval excluding 0.

## RESULTS

Group comparison statistics are presented in Table 1. All groups had fewer women than men, and the between-group gender difference was not significant (χ^2^ (6, N = 97) = 0.59, *p* = 0.997). The LD and LD+DA groups had similar MDS-UPDRS Part III scores both ON and OFF medication, indicating similar levels of motor impairment (OFF: t(44) = 0.68, *p* = 0.50; ON: t(43) = 0.18, p = 0.860). The LD+DA group had a significantly higher levodopa equivalent dose (LEDD) (U = 85.50, *p* <0.001, Table 1) and longer disease duration (t(58) = 3.602, *p* = 0.0007, Table 1) than the LD group.

**Table 1:** Participant characteristics UPDRS III = Unified Parkinson’s Disease Rating Scale, Part III (motor); BDI = Beck’s Depression Inventory; LARS = Lille Apathy Rating Scale; TCI-NS = Temperament and Character Inventory – Novelty Seeking; QUIP-RS = Questionnaire for Impulsive-Compulsive Disorders in Parkinson’s Disease: Rating Scale; PGSI = Problem Gambling Severity Index; MoCA = Montreal Cognitive Assessment, * = p <0.05 compared to Controls or between the two patient groups. The reported data exclude the patient with the MoCA score < 24. ^a^ n=26, data not available for 1 participant.

| **Group** | **Controls**<br>**(n=36)** | **LD (n=34)** | **LD+DA (n=27)** |
| --- | --- | --- | --- |
| Age (SD) | 64.4 (8.24) | 65.4 (8.40) | 65.3 (7.07) |
| Sex: n (%) female | 8 (22.22) | 6 (17.65) | 4 (14.81) |
| Handedness: n (%) left-handed | 2 (5.56) | 6 (17.65) | 1 (3.70) |
| Pharmacotherapy: n (%) |  |  |  |
| Levodopa/ carbidopa IR + CR | - | 14 (41.18) | 12 (44.44) |
| Levodopa/ carbidopa CR | - | 2 (5.88) | 3 (11.11) |
| Levodopa/ carbidopa IR | - | 11 (32.35) | 6 (22.22) |
| Pramipexole | - | - | 23 (85.20) |
| UPDRS III ON (SD) | - | 23.13 (10.95) | 21.44 (10.00) |
| UPDRS III OFF (SD) | - | 23.52 (10.50) | 24.51 (10.52) |
| Levodopa Equivalent Dose<br>(LEDD) (mg/day) (SD) | - | 672.00<br>(334.30) | 1178.00*<br>(345.30) |
| Disease Duration (years) (SD) | - | 4.77 (3.34) | 7.98 (3.53)*<sup>a</sup> |
| Levodopa-Only LEDD (mg/day)<br>(SD) | - | 672.00<br>(334.30) | 821.43 (273.67) |
| BDI (SD) | 6.00 (4.74) | 6.74 (3.90) | 5.56 (3.62) |
| LARS (SD) | -24.61 (6.47) | -26.71 (4.99) | -22.09 (12.09) |
| TCI-NS (SD) | 8.33 (3.25) | 5.60* (3.09) | 6.50 (3.09) |
| QUIP-RS (SD) | 1.15 (1.44) | 0.29* (0.52) | 0.91 (1.13) |
| PGSI (SD) | 0.41 (0.70) | 0.09* (0.30) | 0.22 (0.60) |
| MoCA (SD) | 27.86 (1.61) | 28.06 (1.95) | 27.44 (1.87) |

BDI and LARS scores were comparable among all groups (Table 1), indicating minimal depressive symptoms and apathy (BDI, F(2, 92) = 0.61, *p* = 0.546; LARS, H(3) = 2.85, *p* = 0.240). MoCA scores were also similar among groups (H(3) = 2.10, *p* = 0.350). A Kruskal-Wallis test with Dunn’s multiple comparison procedure revealed a significant difference between groups for QUIP-RS and TCI-NS scores (QUIP, H(3) = 10.08, *p* = 0.007; TCI-NS, H(3) = 13.75, *p* = 0.001). Specifically, participants in the LD group scored significantly lower on both the QUIP-RS (*p* = 0.007) and TCI-NS (*p* < 0.001) than Controls and LD+DA patients, suggesting reduced novelty-seeking and impulsive-compulsive tendencies in the LD group, with no difference between the LD+DA and Controls. There were no significant group differences on the PGSI (H(3) = 5.79, *p* = 0.055), though the LD group showed a non-significant trend towards the lowest score (*p* = 0.058).

### Performance on the VGT

In the Control group, there was an expected risk-promoting effect of Cues (*b* [95% CI] = 0.36 [0.22, 0.50]) (Figure 2A, see supplementary Table S1 for complete model statistics). A significant Cues x EVR interaction (*b* [95% CI] = 0.16 [0.05, 0.26]) indicated that the risk-promoting effect of Cues was value-dependent: cues promoted riskier choice mainly at risk-favoring EVRs. The main effect of EVR was also significant (*b* [95% CI] = 2.86 [2.44, 3.31]).

**Figure 2:**
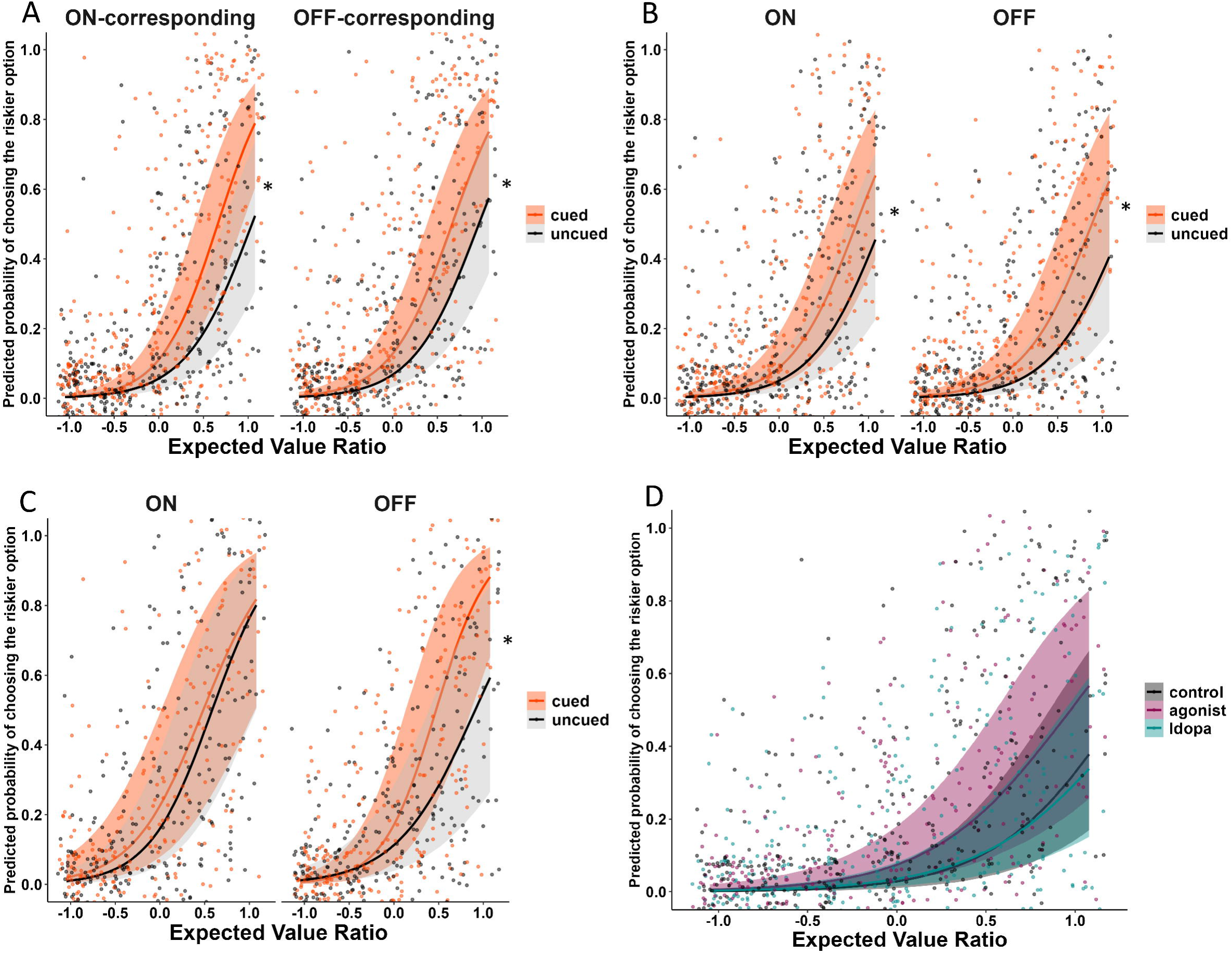
Bayesian model prediction plots depicting predicted likelihood of choosing the riskier option on the Vancouver Gambling Task (VGT). The ribbons represents a 95% Bayesian credible interval. The points represent each participant’s mean likelihood of choosing the riskier option for each participant based on raw data. A) Control group: risky choice on the uncued (grey) and cued (orange) VGT. ON-corresponding = session corresponding to the ON session in the matched patient; OFF-corresponding = session corresponding to the OFF session in the matched patient. The risk-promoting effect of cues is significant. B) Levodopa monotherapy (LD) group: risky choice on the uncued (grey) and cued (orange) VGT ON and OFF medication. The risk-promoting effect of cues is significant. C) Levodopa + DA agonist group (LD+DA): risky choice on the uncued (grey) and cued (orange) VGT ON and OFF medication. The plot depicts a significant Cues x Medication x EVR interaction with the value-dependent effect of cues eliminated in the ON condition. D) Risky choice on the uncued VGT in the OFF condition in the control group (black), the LD group (cyan), and the LD+DA group (deep pink). Group differences are not significant. * = statistically significant finding in the Bayesian models, i.e. the 95% credible interval on a given parameter does not include 0.

In the LD group, participants were also more likely to make risky choices in the presence of cues (*b* [95% CI] = 0.34 [0.18, 0.52]; Figure 2B, Table S2). However, there was no significant effect of Medication (*b* [95% CI] = 0.02 [-0.25, 0.28]) or Cues x Medication interaction (*b* [95% CI] = -0.04 [-0.18, 0.09]). There was also no significant Cues x EVR interaction (*b* [95% CI] = 0.06 [-0.05, 0.17]) demonstrating an absence of a value-dependent effect of Cues. However, the probability direction value of .88 indicated 88% certainty that the risk promoting effect of cues occurred preferentially at risk-favoring EVRs, similar to Controls. The main effect of EVR was significant (*b* [95% CI] = 2.57 [2.13, 3.06]).

In the LD+DA group, there was a significant main effect of Cues (*b* [95% CI] = 0.22 [0.02, 0.43]), a significant two-way Cues x EVR interaction (*b* [95% CI] = 0.19 [0.04, 0.34]), and a significant three-way Cues x Medication x EVR interaction (*b* [95% CI] = -0.34 [-0.51, - 0.18]; Figure 2C, Table S3). The origin of the three-way interaction was that, while Cues significantly and value-dependently increased the choice of riskier alternatives uniquely in the OFF state, performance did not differ on the cued versus the uncued task in the ON state. As in the other groups, there was a significant effect of EVR (*b* [95% CI] = 2.73 [2.02, 3.5]). A post-hoc analysis to further examine the 3-way interaction indicated that medication value-dependently increased risky choice on the uncued task, with choice becoming riskier at more risk-favoring EVRs (Medication x EVR: *b* [95% CI] = 0.38 [0.16, 0.61], Table S4). However, on the cued task, there was a significant Medication x EVR interaction in the opposite direction, with a decrease in value-dependence of choice on medication, i.e. an increase in risky choice at less risk-favoring EVRs and a decrease at more risk-favoring EVRs (*b* [95% CI] = -0.4 [-0.64, - 0.17], Table S5). Thus, LD+DA medication reduced the difference between cued and uncued tasks by encouraging value-dependent risky choice on the uncued task and less value-dependent choice on the cued task.

The three groups did not differ significantly in the likelihood of choosing the riskier option on the uncued task in the OFF-medication state or corresponding session for Controls (Figure 2D, Table S6). Across all groups, there was a significant effect of EVR (*b* [95% CI] = 2.82 [2.28, 3.35]): as expected, participants were more likely to choose riskier options at more risk-favoring EVRs. Additionally, there was a significant effect of Order (*b* [95% CI] = -0.55 [- 0.91, -0.20]), indicating that participants were less likely to make risky choices across task versions and sessions if they were tested OFF medication first.

## DISCUSSION

This study examined individual and combined effects of reward-paired sensory cues and two types of DRT, levodopa monotherapy and combination levodopa with D2/3 receptor agonists, in patients with PD. Cues promoted riskier choice across all groups, mostly in a value-dependent manner. Levodopa alone did not have a significant effect on risk-taking propensity. However, contrary to our hypothesis, combination levodopa and DAs reduced the value-dependent risk-promoting effect of cues in these patients by *making choice riskier in the absence of cues*, while also making choice less value-dependent (i.e. less rational) in the presence of cues. This suggests that LD+DA encourages risk independent of contextual factors, rather than enhancing their effect.

The elimination of the risk-promoting effect of cues under LD+DA indicates sub-additive effects of medication and cues in these patients. In the current study, this sub-additivity may stem from either a) a ceiling effect on the behavioral response or b) one effect functionally limiting the other. A ceiling effect appears unlikely: though the LD+DA group was non-significantly more risk-seeking than the other two groups ‘at baseline’, the patients’ VGT performance did not approach 100% risky choice even at the most risk-favoring EVRs and was more risk-averse than uncued VGT performance of young healthy participants in our previous research (26,27).

More plausibly, the effect of medication may have limited the risk-promoting effect of cues. This may reflect a drug-dependent reduction of dopamine system responsiveness to motivationally salient stimuli against the background of elevated post-synaptic dopaminergic signals. By tonically stimulating post-synaptic D2/3 receptors, DAs may decouple post-synaptic responses from presynaptic dopamine signals, masking the post-synaptic effects of cue-evoked dopamine transients. This could effectively degrade informational content of stimulus-evoked dopamine responses and reduce signal-to-noise ratio. DAs have previously been found to reduce reward prediction error-related activity in healthy controls (50, 51) and in patients with PD (52) and obsessive-compulsive disorder (53). They have also been found to boost salience of both relevant and irrelevant stimuli (54–56). Our observation of reduced value-dependence of choice under LD+DA also aligns with a drug-dependent decrease in signal-to-noise ratio and a flattening of the salience landscape: LD+DA appears to bias individuals towards riskier choices independent of salient reward-paired stimuli or subtle differences in expected values of prospective rewards.

Though the effects of LD+DA appear at odds with rodent findings demonstrating mutually-potentiating effects of D2/3 agonists and cues on risky choice (32,35), chronic ropinirole reversed a positive association between ventral tegmental and nucleus accumbens activity in the cued rodent gambling task (32). This functional decoupling aligns with a drug-dependent untethering of pre- and post-synaptic dopamine signals, proposed above.

A complementary mechanistic explanation is that stimulation of D2 autoreceptors by DAs could downregulate dopamine release from midbrain neurons (57,58). Low DA doses act preferentially at presynaptic striatal D2 receptors to decrease dopamine release (59,60). In rats performing the rodent gambling task, chronic ropinirole decreased c-Fos expression in ventral tegmental dopamine neurons (32). This could produce a state similar to reward deficiency, which is proposed to drive reward-seeking behavior in individuals with substance use disorders (61). Notably, dysfunctional activation of midbrain dopamine autoreceptors has been reported in patients with PD and pathological gambling (62). Together, pre- and post-synaptic effects of DAs could produce a state of increased tonic dopaminergic drive resulting from direct stimulation of post-synaptic D2/3 receptors, coupled with reduced effective gain of phasic signals. This could bias behavior towards globally elevated, cue-insensitive risk-taking. Disordered gambling may emerge from interactions of these acute medication effects with pre-exiting vulnerabilities and/or disease- or therapy-driven chronic alterations in pre- and post-synaptic dopamine signaling, which have been documented in PD patients with pathological gambling (62–65).

The absence of a risk-promoting effect of levodopa monotherapy is surprising given our previous report that levodopa increased risky choice on the VGT in PD (15). This discrepancy could be explained by sample and task characteristics. In the current sample, patients on LD were on higher doses and were more likely to be prescribed the continuous release form. Although continuous release medications were withheld for 18 hours, this washout may have been insufficient, considering long-term treatment and prolonged effective half-life. Indeed, patients in the earlier study were significantly more risk-averse than controls in the OFF state (15), whereas uncued task performance was not significantly different between groups in the OFF state in the current study. The other, less likely explanation are task version differences, as the earlier study employed a version of the VGT, with basic visual cues but no auditory feedback.

### Limitations

Conjectures about the mechanistic underpinnings of the behavioral findings are speculative due to the lack of direct measurement of neural correlates. Furthermore, though the cued and uncued VGT is a sensitive assay of cue-induced risky choice, designed to translate rodent findings to humans (26) it does not lend itself to reinforcement learning modeling and cannot be used to computationally evaluate proposed mechanisms. However, identification of potential theoretical explanations for these findings helps inform future studies.

The dual therapy in the LD+DA group does not permit the evaluation of unique effects of DAs. Because DA monotherapy is uncommon in real-world prescribing practices (66), studying effects of dual therapy offers higher ecological validity. However, the clinical significance of the current work is limited by the absence of patients experiencing (or with a history of) ICDs on dual therapy. In practice, agonist therapy is likely to be withdrawn or minimized for patients who develop ICDs (5,7). It is, therefore, possible that the dual-therapy sample represent a group resilient to ICDs, and the effects of DAs on risky choice may differ in those experiencing ICDs and iatrogenic gambling disorder. Further, given physicians’ cautious approach to prescribing DAs, patients on them may differ clinically and demographically from those on levodopa monotherapy (e.g. greater disease severity), although this was not observed in the current sample. However, an important consideration are the higher LEDDs in the LD+DA group relative to the LD group: besides reflecting the effect of DAs, the findings could in part be driven by the overall higher DRT doses. As mentioned earlier, it is possible that the washout was insufficient to achieve a truly drug-free state. Additionally, “wearing off” may have differed across participants in the ON state due to individual differences, including disease duration (67). Finally, MDS-UPDRS-III could not be blindly rated for some participants.

### Future Directions

The current findings suggest that DAs could produce a state of increased dopaminergic drive, coupled with reduced effective gain of phasic dopaminergic responses to salient stimuli. At the behavioral level, such a state could result in increased motivation to pursue rewards and risk-taking (68–71), decoupled from contextual factors. This broadly aligns with the clinical presentation of ICDs. Future work could investigate neurobiological mechanisms proposed to underpin the behavioral effects of DAs reported here using targeted experimental paradigms in animal models and in humans with or without PD experiencing ICDs on DAs.

### Conclusions

The present study evaluated the effects of levodopa and DAs on cue-induced risky choice in patients with PD. Consistent with previous findings in rats and healthy humans, audiovisual reward cues promoted riskier choice in both patients with PD and age-matched healthy controls. While levodopa did not have a significant effect on risky decision making, a combination of levodopa with DAs eliminated the risk-promoting effect of cues by promoting riskier choice in the *absence* of cues and elevating risk-taking to the level seen with cues in these patients.

Because DAs, combined with levodopa, promote risk-taking even in the absence of reward-paired cues, this combination therapy could translate into risky reward-seeking behaviors decoupled from contextual factors.

## ACKNOWLEDGEMENTS AND DISCLOSURES

The authors thank Carmen Feng for help with performing patient chart reviews.

