## Supplementary tables for "Dopamine Agonists in Parkinson’s Disease Decouple Risk-Taking from Reward-Paired Cues by Enhancing Risky Choice in Their Absence"

**Supplemental Information**

| Parameter | Median [95% CI] | pd | R-hat | ESS |
| --- | --- | --- | --- | --- |
| **Intercept** | **-1.89 [-2.34, -1.44]** | **1.00** | **1** | **2,737.61** |
| **Cues** | **0.36 [0.22, 0.50]** | **1.00** | **1** | **6,819.74** |
| Session | -0.09 [-0.27, 0.08] | 0.86 | 1 | 5,581.99 |
| **EVR** | **2.86 [2.44, 3.31]** | **1.00** | **1** | **1,593.94** |
| Version | -0.1 [-0.21, 0.01] | 0.96 | 1 | 6,878.75 |
| Order | -0.37 [-0.84, 0.09] | 0.94 | 1 | 2,128.74 |
| Trial | -0.03 [-0.14, 0.07] | 0.73 | 1 | 4,277.81 |
| Cues x Session | 0.01 [-0.11, 0.13] | 0.55 | 1 | 6,044.61 |
| **Cues x EVR** | **0.16 [0.05, 0.26]** | **1.00** | **1** | **9,195.35** |
| Session x EVR | 0.07 [-0.03, 0.17] | 0.92 | 1 | 16,709.07 |
| Cues x Session x EVR | 0.07 [-0.03, 0.17] | 0.93 | 1 | 15,157.02 |

Table S1: Statistics for the Bayesian model testing the effects of cues on risky decision making in the Control group. Session reflects repeated testing of the Control group (two sessions): for each Control participant the values of this variable correspond to those of the Medication variable for the matched patient. Significant effects are **bolded.** EVR = expected value ratio; UPDRS III = Movement Disorders Society Unified Parkinson’s Disease Rating Scale Part III (motor); PD = probability direction; ESS = effective sample size. Model formula: *Risky choice ~ Cues*Medication*EVR + Task Version + UPDRS OFF + Order + Trial + (1|id) + (0 +EVR|id) + (0+Trial|id) + (0+Cues*Medication|id)*

| Parameter | Median [95% CI] | pd | R-hat | ESS |
| --- | --- | --- | --- | --- |
| **Intercept** | **-2.24 [-3.05, -1.45]** | **1.00** | **1** | **2,410.88** |
| **Cues** | **0.34 [0.18, 0.52]** | **1.00** | **1** | **4,016.57** |
| Medication | 0.02 [-0.25, 0.28] | 0.56 | 1 | 3,557.16 |
| **EVR** | **2.57 [2.13, 3.06]** | **1.00** | **1** | **1,516.28** |
| UPDRS III OFF | -0.24 [-0.99, 0.53] | 0.73 | 1 | 2,512.82 |
| Version | -0.03 [-0.15, 0.09] | 0.67 | 1 | 5,684.19 |
| Order | -0.21 [-0.77, 0.36] | 0.77 | 1 | 2,512.05 |
| Trial | -0.11 [-0.23, 0] | 0.97 | 1 | 3,804.79 |
| Cues x Medication | -0.04 [-0.18, 0.09] | 0.73 | 1 | 5,090.95 |
| Cues x EVR | 0.06 [-0.05, 0.17] | 0.88 | 1 | 14,241.04 |
| Medication x EVR | 0.05 [-0.06, 0.16] | 0.81 | 1 | 14,538.04 |
| Cues x Medication x EVR | 0.01 [-0.1, 0.12] | 0.57 | 1 | 14,074.59 |

Table S2: Statistics for the Bayesian model testing the effects of cues and medication state on risky decision making in the levodopa (LD) group. Significant effects are **bolded.** EVR = expected value ratio; UPDRS III = Movement Disorders Society Unified Parkinson’s Disease Rating Scale Part III (motor); PD = probability direction; ESS = effective sample size. Model formula: *Risky choice ~ Cues*Medication*EVR + Task Version + UPDRS OFF + Order + Trial + (1|id) + (0 +EVR|id) + (0+Trial|id) + (0+Cues*Medication|id)*

| Parameter | Median [95% CI] | pd | R-hat | ESS |
| --- | --- | --- | --- | --- |
| **Intercept** | **-1.68 [-3, -0.41]** | **0.99** | **1** | **3,072.63** |
| **Cues** | **0.22 [0.02, 0.43]** | **0.98** | **1** | **4,412.07** |
| Medication | 0.19 [-0.01, 0.39] | 0.97 | 1 | 4,783.58 |
| **EVR** | **2.73 [2.02, 3.5]** | **1.00** | **1** | **1,763.00** |
| UPDRS III OFF | 0.42 [-0.91, 1.72] | 0.75 | 1 | 3,347.62 |
| Version | 0.07 [-0.17, 0.32] | 0.72 | 1 | 4,286.79 |
| Order | -0.31 [-1.23, 0.6] | 0.76 | 1 | 3,059.88 |
| Trial | 0.06 [-0.09, 0.2] | 0.80 | 1 | 4,438.06 |
| Cues x Medication | -0.01 [-0.28, 0.22] | 0.54 | 1 | 4,936.60 |
| **Cues x EVR** | **0.19 [0.04, 0.34]** | **0.99** | **1** | **14,229.52** |
| Medication x EVR | -0.06 [-0.22, 0.1] | 0.77 | 1 | 14,327.70 |
| **Cues x Medication x EVR** | **-0.34 [-0.51, -0.18]** | **1.00** | **1** | **15,290.07** |

Table S3: Statistics for the Bayesian model testing the effects of cues and medication state on risky decision making in the levodopa + dopamine agonist (LD+DA) group. Significant effects are **bolded.** EVR = expected value ratio; UPDRS III = Movement Disorders Society Unified Parkinson’s Disease Rating Scale Part III (motor); PD = probability direction; ESS = effective sample size. Model formula: *Risky choice ~ Cues*Medication*EVR + Task Version + UPDRS OFF + Order + Trial + (1|id) + (0 +EVR|id) + (0+Trial|id) + (0+Cues*Medication|id)*

| Parameter | Median [95% CI] | pd | R-hat | ESS |
| --- | --- | --- | --- | --- |
| **Intercept** | **-1.72 [-3.05, -0.46]** | **0.99** | **1** | **4,225.57** |
| Medication | 0.13 [-0.21, 0.49] | 0.79 | 1 | 4,905.38 |
| **EVR** | **2.53 [1.73, 3.34]** | **1.00** | **1** | **2,444.20** |
| UPDRS III OFF | 0.18 [-1.06, 1.54] | 0.62 | 1 | 4,026.68 |
| Version | 0.22 [-0.13, 0.53] | 0.91 | 1 | 4,712.70 |
| Order | -0.48 [-1.37, 0.46] | 0.85 | 1 | 4,115.51 |
| Trial | 0.07 [-0.13, 0.26] | 0.76 | 1 | 5,665.13 |
| **Medication x EVR** | **0.38 [0.16, 0.61]** | **1.00** | **1** | **11,026.99** |

Table S4: Statistics for the post-hoc Bayesian model testing the effects of medication on risky choice on the uncued Vancouver Gambling Task in the levodopa + dopamine agonist (LD+DA) group. Significant effects are **bolded.** EVR = expected value ratio; UPDRS III = Movement Disorders Society Unified Parkinson’s Disease Rating Scale Part III (motor); PD = probability direction; ESS = effective sample size. Model formula: *Risky choice ~ Medication*EVR + Task Version + UPDRS OFF + Order + Trial + (1|id) + (0 +EVR|id) + (0+Trial|id) +(Medication|id)*

| Parameter | Median [95% CI] | pd | Rhat | ESS |
| --- | --- | --- | --- | --- |
| **Intercept** | **-1.74 [-3.14, -0.36]** | **0.99** | **1** | **3,936.94** |
| Medication | 0.17 [-0.16, 0.48] | 0.87 | 1 | 4,870.10 |
| **EVR** | **2.93 [2.02, 3.87]** | **1.00** | **1** | **1,797.57** |
| UPDRS III OFF | 0.73 [-0.69, 2.17] | 0.86 | 1 | 4,465.92 |
| Version | -0.07 [-0.36, 0.21] | 0.70 | 1 | 5,421.20 |
| Order | -0.16 [-1.19, 0.82] | 0.63 | 1 | 4,240.20 |
| Trial | 0.09 [-0.09, 0.24] | 0.86 | 1 | 5,748.85 |
| **Medication x EVR** | **-0.4 [-0.64, -0.17]** | **1.00** | **1** | **9,905.71** |

Table S5: Statistics for the post-hoc Bayesian model testing the effects of medication on risky choice on the cued Vancouver Gambling Task in the levodopa + dopamine agonist (LD+DA) group. Significant effects are **bolded.** EVR = expected value ratio; UPDRS III = Movement Disorders Society Unified Parkinson’s Disease Rating Scale Part III (motor); PD = probability direction; ESS = effective sample size. Model formula: *Risky choice ~ Medication*EVR + Task Version + UPDRS OFF + Order + Trial + (1|id) + (0 +EVR|id) + (0+Trial|id) +(Medication|id)*

| Parameter | Median [95% CI] | pd | R-hat | ESS |
| --- | --- | --- | --- | --- |
| **Intercept** | **-2.71 [-3.48, -1.93]** | **1.00** | **1** | **1,183.26** |
| Group: LD+DA | 0.98 [-0.44, 2.37] | 0.92 | 1 | 1,285.21 |
| Group: LD | 0.22 [-1.06, 1.46] | 0.63 | 1 | 1,126.49 |
| **EVR** | **2.82 [2.28, 3.35]** | **1.00** | **1** | **1,588.78** |
| UPDRS III OFF | -0.43 [-1.04, 0.21] | 0.92 | 1 | 1,289.14 |
| Version | -0.27 [-0.62, 0.08] | 0.94 | 1 | 1,379.46 |
| **Order** | **-0.55 [-0.91, -0.20]** | **1.00** | **1** | **1,374.15** |
| Trial | 0 [-0.15, 0.13] | 0.52 | 1 | 2,509.70 |
| Group: LD+DA x EVR | -0.21 [-1.14, 0.72] | 0.68 | 1 | 1,956.06 |
| Group: LD x EVR | -0.37 [-1.16, 0.4] | 0.82 | 1 | 1,720.48 |

Table S6: Statistics for the Bayesian model comparing the performance of the three groups (Controls, LD group, LD+DA) on the uncued Vancouver Gambling Task in the unmedicated state. Significant effects are **bolded.** EVR = expected value ratio; UPDRS III = Movement Disorders Society Unified Parkinson’s Disease Rating Scale Part III (motor); PD = probability direction; ESS = effective sample size. Model formula: *Risky choice ~ EVR*Group + Task Version + UPDRS OFF + Order + Trial + (1|id) + (0 +EVR|id) + (0+Trial|id)*
